# Diploid origins and early genome stabilization in the allotetraploid *Arabidopsis suecica*

**DOI:** 10.1101/2024.12.06.627142

**Authors:** Robin Burns, Anna Glushkevich, Aboli Kulkarni, Uliana K Kolesnikova, Filip Kolář, Alison Dawn Scott, Polina Yu. Novikova

**Affiliations:** Department of Plant Sciences, University of Cambridge, Cambridge, UK; Department of Chromosome Biology, Max Planck Institute for Plant Breeding Research, Cologne, Germany; Department of Botany, Faculty of Science, Charles University, Prague, Czech Republic; Department of Plant Biotechnology and Bioinformatics, Ghent University, Ghent, Belgium; Flanders Institute for Biotechnology (VIB), VIB-UGent Center for Plant Systems Biology, Belgium

## Abstract

Polyploidization, followed by genome downsizing, is a recurrent evolutionary cycle that dramatically reshapes genome structure. Newly formed polyploids must quickly adjust their cell division machinery to maintain stable chromosome inheritance, while long-term stabilization involves rediploidization, returning the genome to a diploid-like state. Here, we investigate the origin and early genome evolution of *Arabidopsis suecica*, a hybrid polyploid derived from *A. thaliana* and *A. arenosa*

Leveraging a recent genome assembly for *A. suecica* along with population level whole-genome resequencing of all three species, we identify the closest progenitors to *A. suecica* and use this knowledge to examine genes under positive selection and to assess compensatory dynamics between homeologs carrying loss-of-function mutations.

Our findings show that both parental species were diploid, including the paternal *A. arenosa* progenitor. We identify evidence for *de novo* adaptation to allopolyploidy within the *A. arenosa* sub-genome of *A. suecica*, including genes involved in homolog pairing and recombination. Although relaxed purifying selection is evident, likely due to the genome-wide redundancy, we observe functional compensation between homeologous gene pairs in *A. suecica*: when one copy loses function, the other maintains it

Together, these findings revise the origin of *A. suecica* and identify early genome stabilization mechanisms, including evidence for meiotic adaptation and mutational buffering through homeologous gene compensation.

## Introduction

Cycles of polyploidy or recurrent whole-genome duplications (WGD) have profoundly shaped the genomes of most eukaryotes(Aköz & Nordborg, 2019; Hokamp et al., 2003; Van de Peer, 2023). In plants, allopolyploidy, WGD combined with interspecific hybridization, is both widespread(Soltis et al., 1993) and associated with important agronomic traits(Chen et al., 2020; Dubcovsky & Dvorak, 2007; A.-L. Li et al., 2015). Major crops including wheat, cotton, *Brassica*, strawberry, peanut, and tobacco trace their origins to allopolyploid events(Bertioli et al., 2019; Feldman & Levy, 2005; Liston et al., 2014; Renny-Byfield & Wendel, 2014; Sierro et al., 2024; Staudt, 2009; U, Nagaharu, 1935). Understanding the evolutionary success of allopolyploids requires resolving the mechanisms that stabilize their genomes, resolve meiotic incompatibilities, and manage genome-wide genetic redundancy.

The evolutionary fate of allopolyploids depends on the sequence of events that generate them. One key event is whether WGD occurs before or after hybridization because this can influence how readily a stable genome emerges. When WGD follows hybridization, as in *Primula kewensis(Newton & Pellew, 1929), Capsella bursa-pastoris(Bachmann et al., 2021)*, and experimental F₁ hybrids in *Mimulus(Wiiliam & Malcolm, 1971)* and *Arabidopsis(Nasrallah et al., 2000)*, the resulting neo-allotetraploid must adapt *de novo* to coordinate divergent genomes during basic cellular processes. In contrast, if WGD occurs before hybridization through the fusion of 2n gametes, autopolyploid parents can serve as a natural source, since their reduced gametes are equivalent to unreduced gametes from diploids. As a result, the resulting allopolyploid could inherit alleles already adapted to polyploidy, potentially easing genome stabilization. The extent to which inherited alleles are essential, or whether independent adaptations frequently arise in young allopolyploids, remains unclear. This question has important implications for both extinction risk and subsequent allopolyploid genome evolution.

Meiotic stabilization is one of the most immediate and consequential challenges facing newly formed allopolyploids. The formation of multivalents and improper pairing between homeologous chromosomes can disrupt segregation and compromise fertility(de Carvalho et al., 2021; Gonzalo, 2022; Katche, Schierholt, Becker, et al., 2023; Katche, Schierholt, Schiessl, et al., 2023). While divergence between sub-genomes can reduce these risks by limiting homeologous recombination(Ramsey & Schemske, 2002; Soltis et al., 1993; Stebbins, 1947), it does not eliminate it entirely(Gaeta & Chris Pires, 2010; Katche, Schierholt, Becker, et al., 2023; Katche, Schierholt, Schiessl, et al., 2023; Zhang et al., 2020). Experimental allopolyploids, including synthetic *A. suecica*, often exhibit meiotic defects despite a substantial sub-genomic divergence of 11.6%(Burns et al., 2021; Chéron et al., 2023; Nibau et al., 2024).

To achieve stable inheritance, polyploids evolve genetic mechanisms that modify crossover distribution or suppress homeologous recombination. In autopolyploids, this often involves polygenic adaptations in meiotic genes, including genes in the synaptonemal complex(Bohutínská et al., 2021; Bray et al., 2024; Hollister et al., 2012; Yant et al., 2013) and kinetochore(Bray et al., 2024), yet whether these same meiotic pathways are changed in allopolyploids remains an open question. Stable allopolyploids do appear to have evolved genetic control of sub-genome specific recombination. Notable examples include *Ph1* (*ZIP4*)(Martín et al., 2021) and *Ph2* (*MSH7-3D*)(Serra et al., 2021) loci in allohexaploid wheat, the *PrBn* locus in *Brassica napus(Liu et al., 2006)* and *BYS* in natural *A. suecica(Henry et al., 2014)*, all of which suppress homeologous recombination. In yeast hybrids, the mismatch repair pathway has been shown to limit crossovers between divergent chromosomes, and loss of MSH2 can restore fertility in sterile hybrids(Greig et al., 2003). On a transcriptional level, meiotic-related genes were shown to be up-regulated on the *A. thaliana* sub-genome of *A. suecica*(Burns et al., 2021). However, it remains unclear whether meiotic stabilization in young allopolyploids arises primarily through changes in gene expression or protein-coding sequences, or both.

Polyploid genomes do not remain polyploid indefinitely. Ancient WGDs in the ancestry of most diploids suggest that current polyploids will eventually return to a diploid-like state through a process known as rediploidization(Van de Peer et al., 2017). This transition reshapes genome architecture via gene loss, regulatory divergence, and biased fractionation(Baduel et al., 2019; Conover & Wendel, 2022; Mandáková & Lysak, 2018; Monnahan et al., 2019; Redmond et al., 2023; Spoelhof et al., 2021), as seen in both ancient allopolyploids like maize(Sun et al., 2023), and in the diversification of distinct morphotypes in *Brassica*(Cheng et al., 2014). During rediploidization, relaxed purifying selection in redundant gene copies allows deleterious mutations to accumulate, while functional homeologs compensate for their non-functional counterparts, marking early stages of genome downsizing, as observed in *Brachypodium hybridum*(Scarlett et al., 2023). Functional compensation between homeologs may reflect a shift in selective constraint, from individual gene copies to gene pairs, suggesting that polyploid genomes are shaped not just by genetic redundancy, but by selection acting to preserve gene functional integrity. Several studies have shown that deleterious mutations, both point mutations and structural variants, accumulate more rapidly in polyploids than in diploids(Conover & Wendel, 2022; Hämälä et al., 2024; Paape et al., 2018), likely due to masking effects between redundant gene copies. The extent to which this type of mutational buffering between homeologs is shaped by selection, or if it is merely a neutral consequence of genetic redundancy, remains unresolved. Another uncertainty is whether patterns of early gene loss are predictable based on gene function or the sub-genome gene location. While genome dominance and biased fractionation have been reported in some allopolyploid species (Ma et al., 2024; Schnable et al., 2011; Alger & Edger, 2020), they are notably absent in several natural allopolyploids(Burns et al., 2021; Penin et al., 2024; Scarlett et al., 2023; Sun et al., 2023). Gene function has been shown to impact gene fractionation rates, with previous studies showing that genes related to genome integrity and organelle function revert to a single copy level faster than genes involved in signaling, transport, and metabolism, likely due to selective constraints acting on gene dosage levels(Z. Li et al., 2016).

The allotetraploid *A. suecica*, a post-glacial hybrid between *A. thaliana* and *A. arenosa*, is an excellent model for studying the early stages of allopolyploid genome evolution. However the origin of the WGD in *A. suecica*, whether it arose somatically in a diploid hybrid or was inherited via 2x gametes, remains unresolved. If the paternal *A. arenosa* was autotetraploid, WGD likely occurred through the fusion of its reduced gamete (n=2x) with an unreduced *A. thaliana* gamete (2n=2x), a parsimonious route(Jakobsson et al., 2006; Novikova et al., 2017). However, recent evidence from sequence similarity of meiotic genes point to a diploid *A. arenosa* parent(Nibau et al., 2022). A diploid *A. arenosa* parent would imply WGD likely occurred somatically in a diploid hybrid, and that *A. suecica* had to evolve *de novo* solutions to the challenges of polyploidy. Therefore distinguishing between diploid and tetraploid *A. arenosa* as the paternal progenitor is critical for understanding the origin of WGD in *A. suecica*, and for disentangling inherited versus novel mechanisms of meiotic stabilization.

With a high-quality long-read genome assembly of *A. suecica(Burns et al., 2021)* and expanded sequence data from both parental lineages, including the full diversity of *A. arenosa* cytotypes(1001 Genomes Consortium, 2016; Bohutínská, Petříková, Booker, Cobo, et al., 2023b; Burns et al., 2021; Monnahan et al., 2019; Novikova et al., 2016) we address three central hypotheses about the origin and early genome evolution following allopolyploidy in *A. suecica*. First, we ask whether the *A. arenosa* parent of *A. suecica* was diploid or autotetraploid. The paternal *A. arenosa* identity allows us to infer whether WGD likely occurred somatically in a diploid hybrid or was inherited through the hybridization of 2x gametes. Establishing a diploid origin would support the hypothesis that *A. suecica* had to evolve *de novo* solutions to stabilize meiosis and manage genome-wide redundancy. Second, we test whether adaptation to allopolyploidy in *A. suecica* involves positive selection on genes that regulate meiosis and genome integrity. To do this, we identify genes under positive selection in the *A. suecica* genome and use protein-protein interaction networks to determine whether they represent important functional clusters. Finally, we ask how does *A. suecica* manage the mutational load associated with genome-wide genetic redundancy? By analyzing the distribution of inherited and *de novo* loss-of-function (LoF) mutations across sub-genomes, we test for a negative correlation between damaging variants in homeologous gene pairs. In other words we ask, when one copy (e.g., from *A. thaliana*) harbors a LoF mutation, does the other (e.g., from *A. arenosa*) remain intact? Evidence for homeolog compensation would suggest that selection operates at the level of gene pairs, potentially buffering young polyploid genomes against the mutational load that accompanies the polyploid-induced relaxed selective constraint.

## Materials and Methods

### DNA extraction of herbarium samples and sequencing

Herbarium samples were obtained from Moscow University Herbarium(Seregin, 2024). Extraction of DNA from herbarium samples of five *A. arenosa* and one *A. suecica* accession was carried out under a PCR workstation equipped with UV sterilization lights. The workstation, tubes and equipment were UV-sterilized before beginning the extraction. Herbarium samples were weighed and photographed. The material was then crushed using a sterile plastic pestle in a 2ml tube and then weighed again. PTB buffer was added and the material was kept at 37°C overnight on continuous rotation. We extracted the DNA following the protocol of(Gutaker et al., 2017). Libraries were sequenced in 50-bp paired-end mode on Illumina HiSeq 2000 Analyzers at the Vienna BioCenter Core Facilities (VBCF).

### Read mapping and SNP calling

For *A. thaliana* accessions, 970 accessions of *A. thaliana* were used from the 1,001 genomes dataset(1001 Genomes Consortium, 2016), these accessions had a read length greater than 60bp and greater than 8X coverage. In addition to the herbarium samples aforementioned, we used 15 previously published *A. suecica* accessions(Novikova et al., 2017). For *A. arenosa* we used 158 previously published individuals(Bohutínská, Petříková, Booker, & Cobo, 2023; Monnahan et al., 2019) in addition to the 5 herbarium *A. arenosa* individuals sequenced here, giving a total of 163 *A. arenosa* individuals. The *A. arenosa* individuals used had a minimum of 5X coverage. A complete list of the *A. thaliana* and *A. suecica* accessions and the *A. arenosa* individuals used in this study is available as Supplementary File 1.

We mapped illumina paired-end short reads for the 16 *A. suecica* accessions, 970 *A. thaliana* accessions and 163 *A. arenosa* individuals to a chromosome-level genome assembly of *A. suecica*(Burns et al., 2021) using the BWA-MEM algorithm with an increased penalty for unpaired reads set to 15(H. Li & Durbin, 2009), version 0.7.17. The sub-genomes of the *A. suecica* accessions were treated separately after partitioning the mapped short reads. The *A. thaliana* accessions and *A. arenosa* individuals were mapped to the *A. thaliana* and *A. arenosa* sub-genome, respectively. PCR duplicates were removed using the rmdup function from Samtools(H. Li et al., 2009), version 1.10. SNPS were called using Genome Analysis Toolkit (GATK) (McKenna et al., 2010), version 3.8, using the HaplotypeCaller function. Generated gVCFs were combined into a single merged VCF using the combineGVCFs and GenotypeGVCFs functions.

### Variant filtration

We removed multi-allelic sites and indels from our analysis and focused on bi-alleic SNPs. Bi-allelic SNPs were filtered to remove repetitive regions in the *A. suecica* sub-genomes (TEs sequences, centromere repeats, simple repeats and rDNA repeats). SNPs and non-variant sites were removed if the site was less than −0.5x log median coverage or greater than 0.5x log median coverage. The coverage thresholds were chosen to exclude sites with abnormally low or high coverage. Low coverage may indicate poor read mapping or insufficient sequencing depth, while high coverage can signal repetitive elements or cryptic duplications, or potential homeologous exchange events, all of which can lead to erroneous variant calls.

Heterozygous SNPs in *A. thaliana* have previously been shown to often result from segregating gene duplications rather than being residual heterozygosity from outcrossing(Jaegle et al., 2023). To avoid SNP-calling errors from segregating gene duplications in our VCF, we removed heterozygous SNPs that were present in *A. thaliana* and *A. suecica* at a population frequency greater than 5%, since both are highly selfing species. As *A. arenosa* is an obligate outcrossing species, to avoid segregating duplicates we removed heterozygous SNPs that were heterozygous in more than 90% of the individuals in the *A. arenosa* populations we examined. We allowed for 10% missing sites in the *A. arenosa* VCF. We allowed for 15% missing sites on the *A. thaliana* VCF.

The final VCFs contained 9,664,305 biallelic SNPs for the *A. arenosa* sub-genome, and 6,125,591 biallelic SNPs for the *A. thaliana* sub-genome.

### Determining the ploidy level of *A. arenosa* individuals

The cytotype of the published *A. arenosa* individuals ploidy were previously confirmed using FACs(Bohutínská, Petříková, Booker, & Cobo, 2023; Monnahan et al., 2019). nQuire(Weiß et al., 2018) was run on raw fastq files of the 5 herbarium *A. arenosa* individuals, using the ‘denoised’ function and the ‘lrdmodel’. Ploidy of the sample was determined by taking the ploidy that gave the maximum log-likelihood and lowest delta log-likelihood. Ploidy of the herbarium *A. arenosa* individuals was further confirmed by counting the number of different *S*-alleles for each individual with tetraploids having up to four and diploids up to two different *S*-alleles. The S-alleles were identified as described in(Kolesnikova et al., 2023) using short-read data and NGSGenotyp pipeline(Genete et al., 2020) with known *Arabidopsis* SRK sequences as a database.

### Nucleotide divergence analysis

Pairwise nucleotide diversity (π) was calculated using custom scripts (see Code availability) in python. In order to carry out the network based on Nei’s D, we read the VCF into R using the package vcfR(Knaus & Grünwald, 2017) and calculated Nei’D using StAMPP(Pembleton et al., 2013). To visualize the network we used the R package tanggle(Klaus Schliep, Marta Vidal-Garcia, Claudia Solis-Lemus, Leann Biancani, Eren Ada, L. Francisco Henao Diaz, Guangchuang Yu, 2021) and ggtree(Yu, 2020). Geographic maps were made using the ggspatial package in R (Dunnington, 2017)

### Niche modeling of *A. suecica*

We modeled the niche of *A. suecica* using the current occurrence records and the selected Bioclim variables to project it onto the climate at LGM. We used workflow described in the ModleR package in R(Sánchez-Tapia et al., 2020) to the entire modeling (set up, evaluation and projection).

We obtained occurrence data of *A. suecica* from the GBIF database (gbif.org). There are a total of 8,451 records available of which we considered records only from preserved specimens having georeferencing data (1,772 records). This data was supplemented with the published records(Novikova et al., 2017). After cleaning the data for duplicates and occurrences in the same pixel 483 clean data points were obtained. Then, species thinning was carried out using the geo_filt function embdded in ModleR. Finally, 197 data points remained after applying the geographic filter of 0.5 km. These occurrence points were considered for further steps. We created 10000 background points using the setup_sdmdata function in the ModleR package.

We obtained 19 bioclim variables from Worldclim climate data (Fick and Hijmans, 2017) at a resolution of 30 arc seconds. We extracted the data for our occurrence points using the R package raster(Hijmans, 2018). Multicollinearity was checked for the bioclim variables in order to enhance the model consistency and remove the impact of highly correlated variables. Out of the 19 variables, eight uncorrelated variables (bio3, bio4, bio8, bio9, bio10, bio14, bio15) were then used for further analysis

We used maxent to fit our model, which uses the occurrence data and the corresponding climatic data to predict the probability of distribution of the target species. Twenty-five models were created by partitioning the data into five cross-validation partition types and five runs per partition. We evaluated the performance of these models using the true skill statistic (TSS), which is based on the confusion matrix(Allouche et al., 2006). The TSS scores span from 0 to 1, where 0 signifies a model performing no better than chance, and 1 denotes a flawless alignment with the observed data. We had set a TSS value greater than 0.7 to prepare the final model. Since all our models showed TSS values >0.9, we considered all 25 models.

The final model was projected onto the past climate (LGM). Further, occurrence points of *A. thaliana* and *A. arenosa* were overlaid on the projection to understand the origin and distribution of *A. suecica* environmentally and spatially.

### Selection scans

To examine polymorphism patterns in genes we first ran SnpEff for annotation, and we used a modified version of the McDonald Kreitman (MK) test in order to find genes showing signatures of selection in *A. suecica.* We used a modified version of the MK test as *A. suecica* is evolutionarily young (~16 Kya), meaning if we relied on ratios of divergence to polymorphism in the MK test we would often have division by zero. Therefore we focused instead on genes where nonsynonymous divergence (Dn) exceeded synonymous divergence (Ds), nonsynonymous polymorphisms (Pn) were fewer than synonymous polymorphisms (Ps) and where Pn was zero in *A. suecica*.

### Gene ontology (GO) enrichment

We used the R package TopGO(Adrian Alexa, 2017) to perform our GO enrichment analysis, using the ‘weight01’ algorithm to account for the hierarchical structure in GO terms. We kept GO terms with a Fisher test p-value lower than 0.05. We used the gene annotations of *A. thaliana* one-to-one orthologs of *A. suecica* genes. Gene annotations for *A. thaliana* were obtained using the R package biomaRt from Ensembl ‘biomaRt::useMart(biomart = ‘plants_mart’, dataset = ‘athaliana_eg_gene’, host = ‘plants.ensembl.org’).

### SNP annotation, polarization and homeolog analysis

We annotated SNPs using SNPeff (version 5.2). We used 200 of the genetically closest *A. thaliana* accession to polarize annotated SNPs on the *A. arenosa* sub-genome. We used 29 individuals of the closest diploid *A. arenosa* to polarize annotated SNPs on the *A. thaliana* sub-genome. This resulted in 3,500,145 annotated and polarized SNPs for the *A. arenosa* sub-genome and 1,205,789 for the *A. thaliana* sub-genome. 11,812 and 11,983 SNPs, on the *A. arenosa* and *A. thaliana* sub-genome respectively, were annotated as predicted LoF polymorphisms that were later used in our homeolog analysis. Predicted LoF polymorphisms were selected by taking SNPs annotated by SNPeff with a high effect. We excluded SNPs with heterozygosity frequencies exceeding 1% in both *A. thaliana* and *A. suecica* to minimize the risk of bias in the LoF mutation analysis, which could be introduced by segregating gene duplicates leading to false signals of heterozygosity. We filtered for homeolog pairs where in *A. suecica* at least one homeolog in the pair carried a LoF mutation, amounting to 414 genes. In instances where a homeolog carried more than one LoF mutation, we selected the LoF mutation that was at a higher population frequency.

### Permutation analysis

To assess the negative correlation of LoF mutations between the *A. thaliana* and *A. arenosa* sub-genomes in *A. suecica*, we conducted two complementary permutation analyses. First, we compared the observed number of homeologous gene pairs with LoF mutations in both sub-genomes to expectations from random pairings. Specifically, we focused on 414 homeolog pairs in which at least one gene copy carried a LoF polymorphism in *A. suecica*. These 414 homeolog pairs represent the subset of the genome where LoF variation exists and where compensatory dynamics can be meaningfully assessed. We did not include all homeologs in the permutation, as the rarity of LoF mutations would heavily skew the null distribution toward zero co-occurrence, limiting the sensitivity of the test. From this filtered set, we generated 1,000 permutations by randomly re-pairing homeologs, and compared the observed number of dual LoF pairs to the resulting distribution.

Secondly, we performed a permutation analysis using simulated *in-silico A. suecica* genomes. We generated 1,000 in-silico *A. suecica* by randomly combining genomes from 200 *A. thaliana* and 29 diploid *A. arenosa* individuals that are genetically closest to natural *A. suecica*, sampling with replacement to account for all possible combinations. For each simulated genome, we assessed dual LoF occurrence in the same set of 414 homeolog pairs as in the natural *A. suecica* dataset. The observed number of homeolog pairs with LoF mutations in both sub-genomes was then compared to the distribution derived from the *in-silico* genomes, providing an additional null expectation under neutral inheritance.

For both tests, statistical significance was determined by comparing the observed number of dual LoF pairs in natural *A. suecica* to the empirical null distribution generated by the permutation.

### Purifying selection analysis

To evaluate evidence of purifying selection acting on intact homeologs relative to their LoF carrying counterpart, we quantified pairwise nucleotide diversity (π) using coding sequences from *A. suecica*. We further calculated the ratio of non-synonymous (Pn) to synonymous (Ps) polymorphisms to assess selective constraint. Given the severe population bottleneck in *A. suecica*, a pseudocount of 0.1 was added to both Pn and Ps to avoid division by zero. Statistical significance was determined using a one-sided paired t-test, testing the hypothesis that intact homeologs have reduced π and Pn/Ps compared to their LoF pairs.

### Analysis of cell cycle genes in the *A. suecica A. arenosa* sub-genome

Biallelic SNPs from CDS regions of the genes were used to generate SNP-frequency heatmaps. *A.arenosa* assemblies from(Bohutínská, Petříková, Booker, Cobo, et al., 2023a) were scaffolded with RagTag(Alonge et al., 2022) on the *A.lyrata* NT1 genome assembly(Kolesnikova et al., 2023) and annotated with Helixer(Stiehler et al., 2021). *A. thaliana* protein sequences of target genes were used for protein BLAST(Camacho et al., 2009) against annotated proteins extracted with AGAT(Dainat et al., 2024). Genes identified by BLAST were aligned with L-INS-i algorithm of MAFFT(Katoh et al., 2005) and trees were constructed with IQTree(Hoang et al., 2018; Nguyen et al., 2015) with fast bootstrap option equals 1000. Protein alignments were visualized using snipit(O’Toole et al., 2024)

### Single and multi copy gene analysis

Single copy and multi copy genes were obtained from(Z. Li et al., 2016) and hypergeometric tests were performed to test for enrichment.

### Expression analysis

To test whether transcription is biased toward the intact member of a homeologous pair, we mined log₂-fold-change (log₂FC) estimates for rosette tissue from(Burns et al., 2021) which include 15 natural *A. suecica* accessions and two synthetic lines. We first averaged log₂FC values within each group (natural vs. synthetic). We then restricted the analysis to homeolog pairs in which the LoF allele segregates at high frequency in natural *A. suecica* (≥ 60 %). This filter retained 64 intact homeologs, 16 on the *A. arenosa*-derived sub-genome and 48 on the *A. thaliana*-derived sub-genome, out of the 414 total pairs.

### STRING Network Analysis

To investigate protein-protein association networks among genes identified in our selection scan of the *A. arenosa* sub-genome of *A. suecica*, we used the STRING database(Szklarczyk et al., 2023). To identify gene groups with shared function and protein-protein associations, we applied Markov Cluster Algorithm (MCL) clustering, using an inflation parameter of 2, resulting in 102 discrete clusters. Four of the 102 clusters were investigated further due to their association with important polyploid traits.

## Results

### Genetic origins of *A. suecica*

To reconstruct the evolutionary history of allotetraploid *A. suecica*, we first aimed to identify the closest parental accessions, using available data from the parental species, *A. thaliana* and *A. arenosa*, covering their vast geographic range. We mapped 970 *A. thaliana* accessions(1001 Genomes Consortium, 2016) and 163 *A. arenosa* individuals (93 tetraploids and 70 diploids), representing spatially close lineages of both diploid and autotetraploid cytotypes(Bohutínská et al., 2024; Monnahan et al., 2019; Novikova et al., 2016), to the *A. thaliana* and *A. arenosa* sub-genomes of *A. suecica*(Burns et al., 2021). Genomic positions were conservatively filtered to ensure reliable variant calling (see Materials and Methods). In summary, callable sites included ~62% (73,813,230) of the *A. thaliana* sub-genome and ~34% (47,950,105) of the *A. arenosa* sub-genome, with 6,125,591 biallelic single nucleotide polymorphisms (SNPs) in *A. thaliana* and 9,664,305 in *A. arenosa.* Throughout this study, results generated for the sub-genomes of *A. suecica* are compared with corresponding data from the parental species, *A. thaliana* and *A. arenosa*.

Previously, it was shown that the closest *A. thaliana* accessions to *A. suecica* are found in Central Eurasia(Novikova et al., 2017). Here, eliminating the possibility of a reference bias artifact, we calculated median pairwise diversity (π) for each *A. thaliana* accession to our 16 *A. suecica* natural accessions and confirmed that result (Figure 1A). To identify the *A. arenosa* individuals most closely related to *A. suecica* and to determine their cytotype (diploid or tetraploid), we expanded the previously released *A. arenosa* dataset(Bohutínská et al., 2024; Monnahan et al., 2019). We sequenced five additional autotetraploid *A. arenosa* individuals sampled from herbarium specimens collected in the southern Urals, representing the easternmost populations and those geographically closest to the parental *A. thaliana* accessions. The southern Ural *A. arenosa* individuals were identified as tetraploids, using the software nQuire(Weiß et al., 2018) and counting the number of different *S*-alleles (see Materials and Methods, similar to ploidy determination in *A. lyrata*[66]), while the ploidy of most other *A. arenosa* samples was previously determined using flow cytometry(Bohutínská et al., 2024; Monnahan et al., 2019).

**Figure 1:**
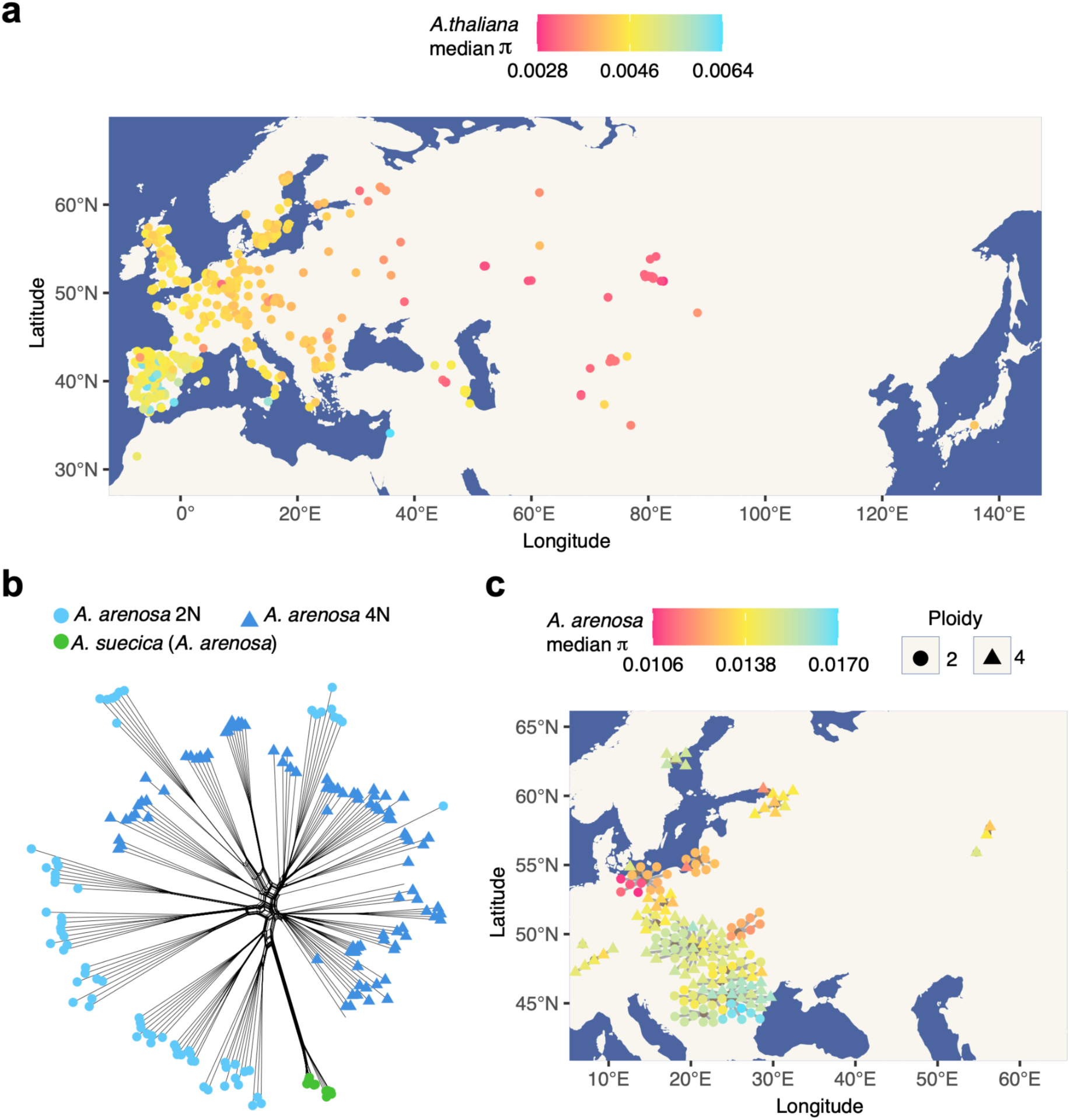
The genetic and geographic origin of *A. suecica*. **a** Median nucleotide diversity (π) of 970 *A. thaliana* accessions to 16 *A. suecica* accessions shows Central Eurasian *A. thaliana* are genetically closest **b** A genetic distance network (Nei’s D) shows that *A. suecica* (green circles) is genetically closer to diploid *A. arenosa* (light blue circles) than autotetraploid *A. arenosa* (dark blue triangles). **c** Median nucleotide diversity (π) of *A. arenosa* individuals to 16 *A. suecica* accessions indicates that Southern Baltic *A. arenosa* diploids (red circles) are the genetically closest. Locations are represented by grey lines connecting *A. arenosa* individuals to their geographic origins.

We conducted a network analysis based on genetic distances (Nei’s D(Nei, 1972)) that are suitable for mixed-ploidy datasets(Pembleton et al., 2013) (Figure 1B). We also calculated the median pairwise nucleotide diversity (π) for each *A. arenosa* individual against our 16 *A. suecica* natural accessions and placed it on a geographical map (Figure 1C). Our results indicate that diploid *A. arenosa* from southern Baltic sea coast(Kolář et al., 2016) is genetically the closest to *A. suecica*, and certainly closer than any autotetraploid *A. arenosa*, including both the current sympatric tetraploid Ruderal lineage growing in Fennoscandia(Monnahan et al., 2019) and tetraploid populations from the southern Urals, i.e. the geographic region closest to putative ancestral *A. thaliana* lineage (Figure 1A). A full list of *A. thaliana* and *A. arenosa* accessions and their π values to *A. suecica* is available in Supplementary File 1. The vast majority of the SNPs (>99%) in *A. suecica* are shared with *A. arenosa* and *A. thaliana,* indicating that polymorphisms in *A. suecica* was largely inherited from standing variation in multiple, genetically distinct parental individuals, rather than arising from *de novo* mutation, in agreement with previous reports(Novikova et al., 2017) (Supplementary Figure 1).

Together our results suggest a diploid rather than tetraploid origin for the *A. arenosa* sub-genome within *A. suecica*. This is in agreement with data showing that meiotic genes on the *A. arenosa* sub-genome of *A. suecic*a carry sequence polymorphisms characteristic of diploid *A. arenosa*, and not the polymorphisms found in autotetraploid *A. arenosa* (Nibau et al., 2022). The diploid origin supports the original hypothesis by Hylander in 1957(Hylander, 1957) that *A. suecica* originated from hybridization between diploid *A. thaliana* and diploid *A. arenosa*, with autotetraploid *A. arenosa* later spreading across Fennoscandia, while the diploids remain restricted to southeastern Europe and isolated Baltic areas(Kolář et al., 2016). The lack of geographical overlap between the current distribution of the closest *A. thaliana* and *A. arenosa* progenitors raises the question of where the hybridisation could have occurred? To resolve this question we performed ecological niche modelling in the context of the estimated time of origin of *A. suecica*, that is, towards the end of the Last Glacial Maximum(Novikova et al., 2017), and we suggest that hybridization between *A. thaliana* and *A. arenosa* likely happened close to the border of the Fennoscandia ice sheet (see Supplementary Note 1, Supplementary Figure 7).

### Genomic adaptation to allopolyploidy in *A. suecica*

Given that the genetically closest parents of *A. suecica* were diploids, *A. suecica* could not inherit any of the potentially pre-adaptive meiotic alleles from autotetraploid *A. arenosa*. Previously, we reported that meiosis-related genes on the *A. thaliana* sub-genome of *A. suecica* are up-regulated compared to parental transcriptional levels(Burns et al., 2021). However, in the same study, we did not observe significant upregulation of meiosis-related genes on the *A. arenosa* sub-genome. Notably, the latter transcriptional comparison was made against tetraploid *A. arenosa*, the previously hypothesized parent of *A. suecica*, whereas we now establish here that diploid *A. arenosa* is closer. Therefore, to uncover how *A. suecica* adapted to allopolyploidy from diploid ancestry, we searched for signatures of selection on genes, performing McDonald-Kreitman test between *A. suecica* and the identified closest progenitors (see Methods). While this approach highlights strong candidates for selection on coding sequences, we note that it will miss selection acting on regulatory elements. Homeologous recombination events could also inflate divergence estimates at some loci; however, we applied coverage-based filters during SNP calling to avoid such events (see Materials and Methods). Additionally, HE events are rare in natural *A. suecica*, with only two homoeologous exchange junctions reported in (Burns et al., 2021), and nine in (Zhang et al., 2020), in a population of *A. suecica*. HE events are also distally biased towards subtelomeric regions(Nibau et al., 2024; Zhang et al., 2020). In contrast, our selection-scan genes do not show such a bias (see Supplemental Figure 2), indicating that HEs are unlikely to drive our selection-scan results. Additionally, gene conversion rates in *A. thaliana* are similarly low, estimated at ~3.6 × 10⁻⁶ per site per meiosis (Wijnker et al., 2013).

Using the MK test, we identified signatures of selection on 35 genes in *A. thaliana* and 585 genes in the *A. arenosa* sub-genomes of *A. suecica* (Supplementary File 2). Due to the low number of founders contributing to the origin of *A. suecica*(Novikova et al., 2017), regions of the genome are depleted for genetic diversity, however, we find that many of the genes under selection are located outside of these low-diversity regions (Supplementary Figure 2), and show GO enrichment, supporting their biological relevance. Enriched categories include meiotic cell cycle (GO:0061780) and mitotic cohesin loading (GO:0051321); however, some are driven by individual genes (Supplementary Figure 3, Supplementary File 2), for example, SCC2 alone drives the mitotic cohesin loading term and so this single-gene driven GO terms limit a broader interpretation. We note that genes under selection on the *A. arenosa* sub-genome do not overlap with the previously described BYS QTL locus for meiotic stability in *A. suecica*(Henry et al., 2014).

To better explore how positively selected genes might functionally interact, we used the STRING database(Szklarczyk et al., 2023) to construct a protein-protein association network (Figure 2A; see Methods). Among the 585 genes under selection in the *A. arenosa* sub-genome of *A. suecica*, 19 formed a connected subnetwork in STRING. These genes clustered into four distinct functional clusters identified using the Markov Cluster Algorithm (MCL), corresponding to sister chromatid cohesion, DNA methylation-dependent heterochromatin assembly, mismatch repair, and spindle elongation. The sister chromatid cohesion cluster contains 6 interacting proteins, including *SCC2*, *WAPL2*, and *SHUGOSHIN*. The DNA methylation cluster includes 4 genes, among them *SUVH2* and *DDM1*. The mismatch repair cluster (5 genes) contains *MutS* and *PMS1* and the spindle elongation cluster (4 genes) includes *MPS1-2.* The four functional clusters are supported by multiple protein-protein association evidence types, including experimentally determined, co-expression, protein homology, and curated databases (Figure 2A legend). In contrast, the 35 selected genes from the *A. thaliana* sub-genome did not form a connected network or enriched functional clusters.

**Figure 2:**
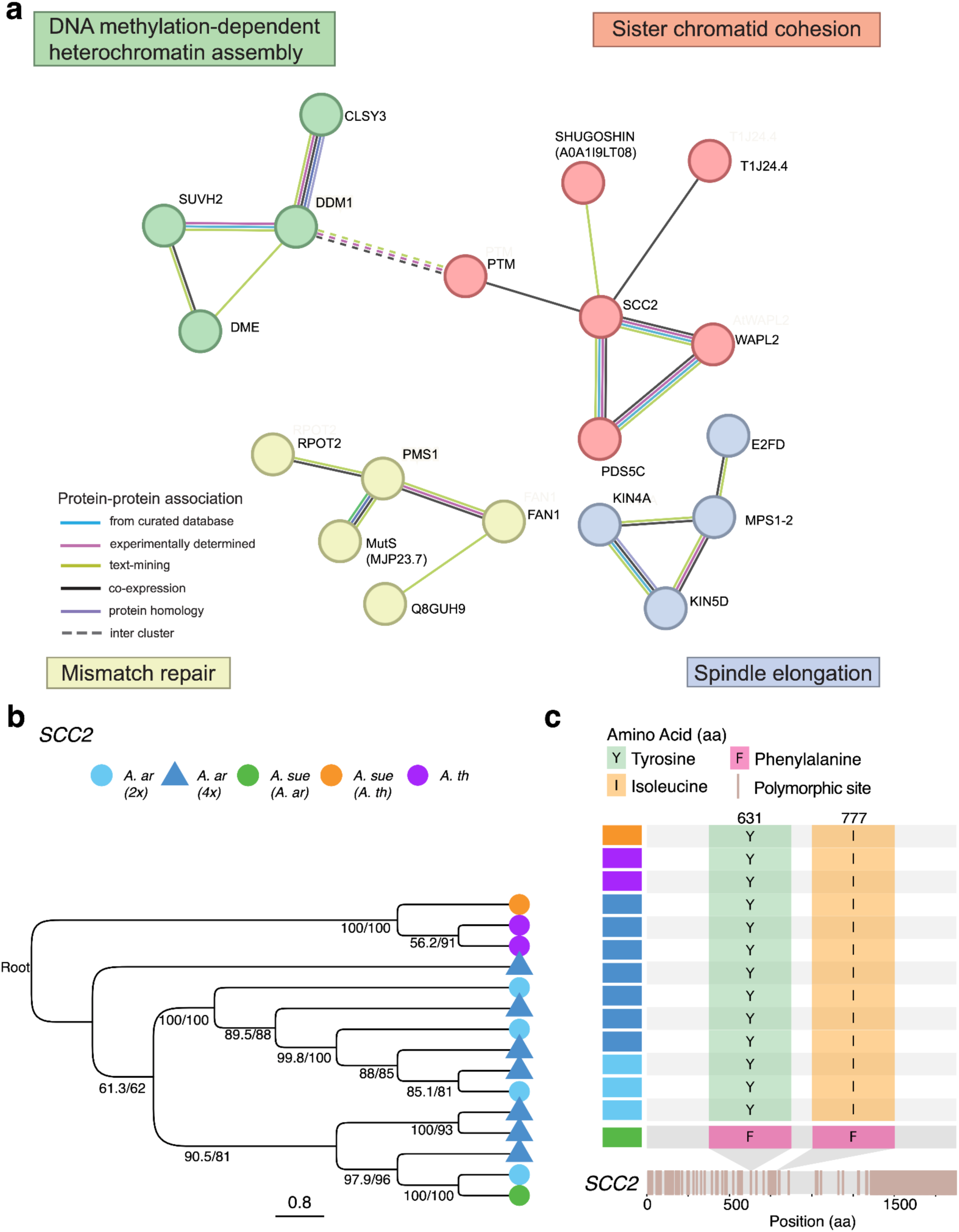
Genomic adaptation to allopolyploidy in *A. suecica*. **a** STRING protein-protein association network showing four functional clusters among 19 genes under positive selection on the *A. arenosa* sub-genome of *A. suecica*, as identified by McDonald–Kreitman tests. Colored clusters represent functional clusters including DNA methylation-dependent heterochromatin assembly (green), sister chromatid cohesion (red), mismatch repair (yellow), and spindle elongation (blue). Solid lines represent intra-cluster associations; dotted lines indicate inter-cluster associations **b** Phylogenetic tree from protein alignment of SCC2 showing that the *A. arenosa* homeolog in *A. suecica* clusters with diploid *A. arenosa*. Branch length is not drawn to scale to improve the analysis of the tree topology. Both the translated genomic and CDS sequence of the *SCC2* from *A. thaliana,* in addition to the *A. suecica A. thaliana* homeolog were used to root the tree. **b** Fixed amino acid differences in the *SCC2* between the *A. suecica A. arenosa* sub-genome and diploid *A. arenosa*. Tetraploid *A. arenosa* and *A. thaliana* share the same amino acid as diploid *A. arenosa*, the difference is unique to *A. suecica*.

Among the four clusters, the sister chromatid cohesion cluster stood out because of the importance of cohesion and the cohesin complex in meiosis(H. Wang et al., 2020)(Storchová et al., 2006). *SCC2* (SISTER CHROMATID COHESION2), which encodes the cohesin loader adherin, a protein essential for cohesin loading onto chromosomes(H. Wang et al., 2020), emerged as a particularly compelling candidate because *SCC2* has also previously been implicated in adaptation in autotetraploid *Arabidopsis arenosa*(Yant et al., 2013). To investigate whether SCC2 represents the same haplotype as that of tetraploid *A. arenosa* or instead represents *de novo* evolution within *A. suecica* following allopolyploidization, we performed a phylogenetic analysis which showed that *A. suecica* clusters consistently with diploid *A. arenosa*(Figure 2B), and does not show evidence of haplotype sharing with autotetraploid *A. arenosa*. Furthermore, *A. suecica* has private polymorphisms on SCC2 (*A. arenosa* sub-genome), which are not present in either diploid or tetraploid *A. arenosa* or *A. thaliana* (Figure 2C), suggesting *de novo* evolution.

### Preservation of gene function via homeolog compensation in *A. suecica*

Polyploid organisms have a substantial amount of genetic redundancy, which can lead to relaxed purifying selection. This pattern of relaxed selection has been observed in autopolyploid *A. arenosa*(Baduel et al., 2019)(Vlček et al., 2025) and allopolyploid *A. kamchatica*(Paape et al., 2018). We investigated whether *A. suecica*, the youngest polyploid in the *Arabidopsis* genus, shows a similar signal of relaxed purifying selection and asked how patterns of LoF mutations are distributed across its sub-genomes.

Analyzing over 3.5 million SNPs in the *A. arenosa* sub-genome and 1.2 million SNPs in the *A. thaliana* sub-genome, we found that both sub-genomes in *A. suecica* are evolving under purifying selection, however purifying selection appears relaxed in *A. suecica* compared to the parental species (Figure 3A). This genome-wide signal suggests a reduced selective constraint following polyploidization, consistent with the buffering effects of homeolog redundancy.

**Figure 3:**
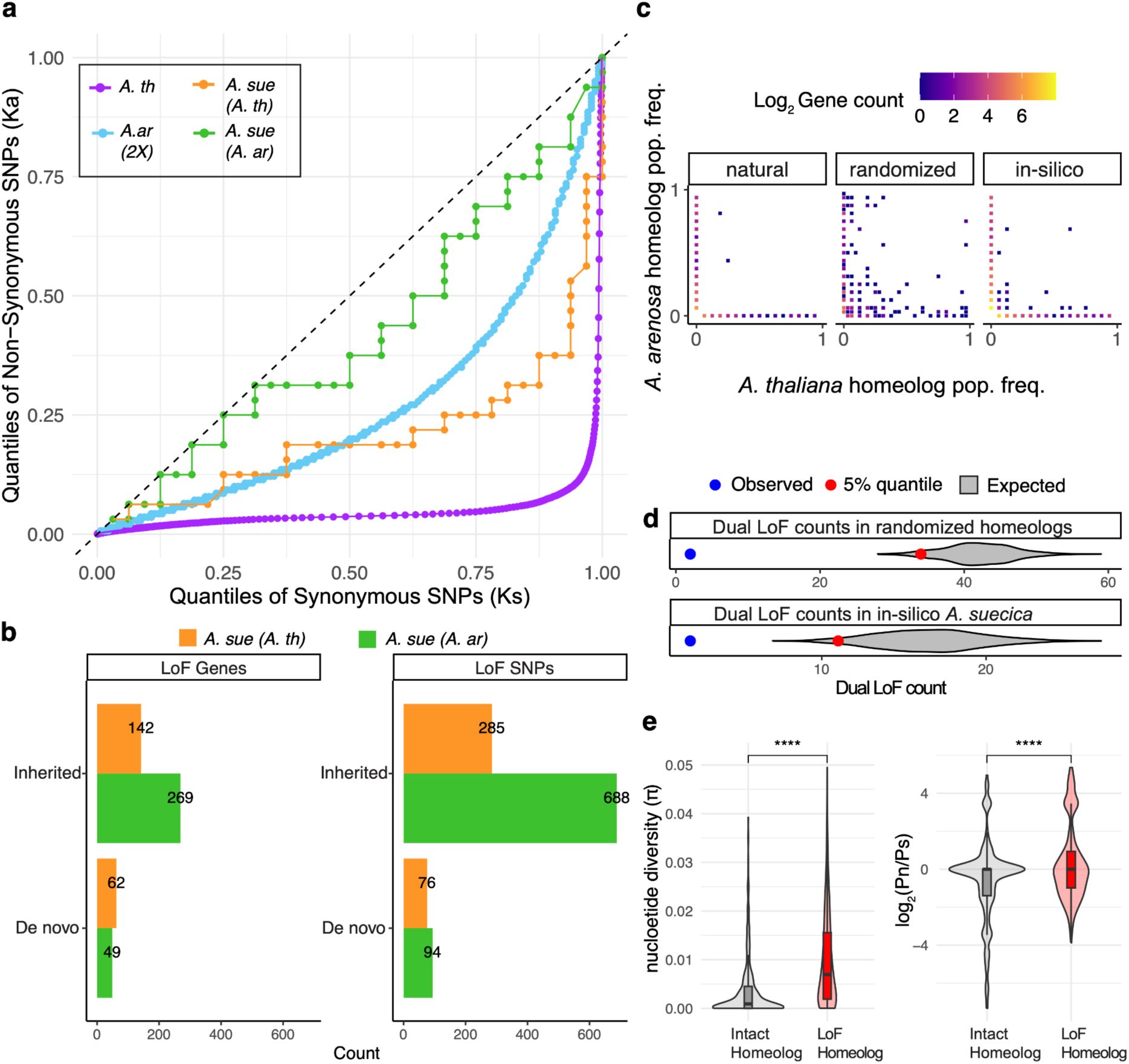
Relaxed purifying selection and homeolog compensation in *A. suecica*. **a** QQ plot of synonymous (Ks) and non-synonymous (Ka) SNP frequencies for *A. suecica* and its parent species (*A. thaliana* and diploid *A. arenosa*).The dotted line indicates neutral evolution. All species exhibit purifying selection, with the *A. suecica* sub-genomes showing relaxed purifying selection relative to the parents. **b** Bar plots showing the number of genes and total counts of LoF SNPs in *A. suecica*, the majority of which are inherited from the parent species. **c** Joint frequency spectra of homeolog pairs with at least one LoF SNP, displaying population frequencies for *A. thaliana* and *A. arenosa* homeologs on the x- and y-axes, respectively. From left to right: observed spectrum in natural *A. suecica*, example from permutations of randomized homeologs and in silico *A. suecica* controls (random combinations of *A. thaliana* and *A. arenosa* individuals). The number of homeologs analyzed are the same (414 homeologs), but the distribution of LoF mutations are different between the three panels. **d** Comparison of natural *A. suecica* with 1,000 randomized and in-silico controls, showing that *A. suecica* (blue dot) has fewer dual LoF SNPs in homeolog pairs than expected by chance (gray violin plot), a difference that is statistically significant (red dot denotes p-value = 0.05) **e** Evidence of purifying selection acting on the intact homeolog, which shows significantly lower nucleotide diversity (π) and a lower ratio of non-synonymous (Pn) to synonymous polymorphisms (Ps) than the LoF homeolog, within *A. suecica.* Statistical significance is indicated by stars (one-sided *t*-test).

A reduced selective constraint following polyploidization enables mutational robustness between the sub-genomes, since a LoF mutation in one sub-genome can be compensated by its homeolog on the alternate sub-genome. To investigate the compensatory relationship between homeologs we identified a total 414 homeologous gene pairs where at least one copy carries a segregating LoF mutation in *A. suecica*. To understand the nature of LoF mutations between homeologs we split them into inherited (standing variation) and *de novo* (occurring post-allopolyploidization) LoF mutations in *A. suecica*. While a slight excess of LoFs were found in *A. arenosa*-derived homeologs (269 vs. 142), the majority of these mutations are also present in the parental genomes and thus appear to be inherited rather than arising *de novo* (Figure 3B). Focusing only on *de novo* LoF mutations unique to *A. suecica*, the distribution appears more balanced (49 in *A. arenosa*, 62 in *A. thaliana*). To investigate if these LoF homeologs overlap with genes identified in our selection scans for genomic adaptation to allopolyploidy in *A. suecic*a (see previous section), we compared the genes involved. However, we find minimal overlap between the gene lists (Supplementary Figure 4), indicating that a LoF occurring on one homeolog is not driving the signals of positive selection we observed in the previous section.

We next asked if LoF mutations (both inherited and *de novo)* observed in 414 homeologous gene pairs are distributed randomly or whether having a LoF mutation in one homeolog would interfere with occurrence of LoF in the other homeolog. The latter would be explained by purifying selection acting to preserve at least one functional copy in *A. suecica*. To test this, we compared the observed joint frequency spectrum of LoF mutations between the *A. suecica* sub-genomes to (1) spectra resulting from reshuffling of homeolog pairings and (2) to spectra of in-silico synthetic *A. suecica* genomes generated by combining natural diploid *A. arenosa* and *A. thaliana* individuals (Figure 3D). In both comparisons, we found that dual LoF mutations are significantly less frequent in natural *A. suecica* (Figure 3C–D; *p* < 0.05). The depletion of dual LoFs supports a model of compensatory homeolog retention, in which functional redundancy between homeologs allows relaxed selection at the level of individual genes, while purifying selection operates at the pair level to prevent the complete loss of gene function. To further test for evidence of purifying selection acting on the remaining intact homeolog, we compared pairwise nucleotide diversity (π) and the ratio of non-synonymous to synonymous polymorphisms (Pn/Ps) between our 414 homeolog pairs. We found that intact homeologs exhibited significantly lower π and Pn/Ps values than the paired homeolog carrying a LoF mutation, consistent with purifying selection acting on the intact copy (*p* = 2.14 × 10⁻²⁶ for π; *p* = 5.15 × 10⁻¹⁰ for Pn/Ps; one-sided *t*-tests Figure 3E). Supporting this further, we also observed sub-genome-biased expression favoring the intact copy, a pattern absent in synthetic *A. suecica*, suggesting it is an evolved rather than inherited expression change (Supplementary Fig. 6). (Supplementary Figure 6). We observed no consistent bias in retention from either sub-genome, suggesting that functional constraint, rather than sub-genome dominance, is the primary force shaping homeolog retention in *A. suecica*.

Genes that consistently revert to a single-copy state following whole-genome duplication are thought to be under strong evolutionary constraint, likely because they function optimally at single-copy dosage and are sensitive to the effects of gene duplication(Edger & Pires, 2009; Makino & McLysaght, 2010). To test whether such dosage constraints influence homeolog retention in *A. suecica*, we asked whether genes previously identified as evolving toward single-copy status across angiosperms(Z. Li et al., 2016) are overrepresented among homeologs carrying LoF mutations. We also tested whether LoF mutations in these single-copy genes occur at higher frequencies in *A. suecica* populations compared to those in genes typically retained in multiple copies. Using the published gene set from(Z. Li et al., 2016), we observed no significant enrichment of single-copy genes among homeologs carrying a LoF mutation, nor do we see that LoF mutations within single copy genes exist at higher population frequencies than LoF mutations within multi-copy genes in *A. suecica* (Supplementary Figure 5). These results suggest that constraint on single-copy gene retention is not a major factor shaping homeolog loss of function in *A. suecica*, at least at this stage of genome evolution.

In the three *A. suecica* homeolog gene pairs where we do observe segregating dual LOF mutations, we tested whether functionally redundant copies from older whole-genome duplications (WGDs) might compensate for complete gene function loss. Using ancient homeolog pairs from *A. thaliana* (Guo et al., 2013), we found no overlap with our dual LOF genes, suggesting these losses are not buffered by retained duplicates from older WGDs. The low frequency of dual LOF mutations (3 out of 414 gene pairs) supports the action of strong purifying selection in the population against complete loss of the gene function.

## Discussion

### A diploid origin for *A. suecica*

*A. suecica* is a relatively young allopolyploid species that originated from the hybridization of *A. thaliana* and *A. arenosa*. Using population-level DNA sequencing for all three species, we revisited the genetic origins of *A. suecica* to understand the nature of WGD leading to the formation of the allopolyploid species and if the ancestry shaped adaptation to allopolyploidy.

Leveraging the recent genome assembly of *A. suecica*(Burns et al., 2021), we minimized reference bias and identified the genetically closest individuals of both progenitor species. Consistent with earlier reports(Novikova et al., 2017), we confirm that the *A. thaliana* parent was diploid. However, we now find that the *A. arenosa* lineage closest to *A. suecica* is also diploid, not tetraploid. Together, this supports a model in which *A. suecica* originated via somatic genome doubling in a diploid hybrid background, rather than through fusion of unreduced gametes. A similar pattern is also seen in *A. kamchatica*, another allopolyploid *Arabidopsis* species, which also derives from diploid progenitors(Kolesnikova et al., 2023; Scott et al., 2024). These findings suggest that somatic doubling of diploid hybrids may be a general mechanism for allopolyploid formation in *Arabidopsis*.

### Evolution of meiosis and genome stability in *A. suecica*

Our STRING network revealed four functional clusters, sister chromatid cohesion, DNA methylation, spindle elongation, and mismatch repair that together suggest genomic adaptation to the challenges of recombination and chromosome segregation in allopolyploid meiosis.

The sister chromatid cohesion cluster includes *SCC2* and *WAPL2*, which are central regulators of cohesin dynamics. *SCC2* encodes the conserved cohesin loader required for deposition of cohesin complexes along chromosomes(H. Wang et al., 2020). In *A. thaliana*, SCC2 localizes to meiotic chromosome axes and is essential for axis formation, synapsis, and homolog pairing(H. Wang et al., 2020). *WAPL2*, conversely, promotes cohesin removal, allowing for the timely release of cohesion during meiotic prophase I(Srinivasan et al., 2019; H. Wang et al., 2020) and in *A. thaliana*, loss of *WAPL* leads to unresolved bivalents and uneven chromosome segregation(De et al., 2014). *Shugoshin* has an important role in protecting cohesion at centromeres at anaphase I(Watanabe & Kitajima, 2005). Together these genes support the functional importance of cohesion and the cohesin complex in polyploid meiosis. We note that SCC2 shows independent signatures of selection in both *A. arenosa* and *A. suecica*, suggesting its general importance in relation to polyploid genome stabilization.

The DNA methylation cluster contains *SUVH2* and *DDM1*. *SUVH2* has been previously shown to play an important role in suppressing the transcription of genes that are associated with meiotic double strand breaks (DSBs) (C. Wang et al., 2022) while a deficiency in *DDM1* has been shown to lead to an increase in meiotic crossovers in *A. thaliana*(Melamed-Bessudo & Levy, 2012). Their selection in *A. suecica* suggests the importance of chromatin in allopolyploids possibly by restricting DSBs and recombination to appropriate genomic regions.

The mismatch repair cluster contains the key genes *MutS* and *PMS1*. In allohexaploid wheat, the *Ph2* locus encodes a *MutS* homolog *MSH7-3D* which has been shown to suppress homeologous recombination by destabilizing homeologous interactions (Serra et al., 2021). *MSH7-3D* is thought to influence recombination partner choice and favor homologous over homeologous chromosomes by destabilizing recombination intermediates that contain mismatches(Serra et al., 2021). In yeast, *PMS1* has been shown to reduce recombination between homeolog chromosomes in meiosis(Chambers et al., 2023) and loss of MMR activity in *A. thaliana* has been shown to increase homeologous recombination nine-fold(L. Li et al., 2006).

The spindle elongation cluster which includes the cell cycle kinase *MPS1-2*. *MPS1-2* is essential for spindle pole body duplication and the spindle assembly checkpoint(Jiang et al., 2009), and polyploid yeast have been shown to be critically dependent on *MPS1-2* for successful chromosome segregation(Meyer et al., 2023) suggesting a conserved role in supporting meiotic stability in polyploid genomes.

Taken together, these four clusters suggest a coordinated mechanism by which *A. suecica* has adapted its meiotic machinery to maintain genome stability following allopolyploidization. First, cohesion regulators and cohesin-associated proteins, including *SCC2*, *WAPL2*, and *Shugoshin*, help establish the meiotic axis and form the physical scaffold necessary for proper homolog alignment. Second, DNA methylation and heterochromatin-associated proteins, such as *DDM1* and *SUVH2*, contribute to shaping the chromatin landscape to regulate the formation of DSBs, which are essential for recombination. Third, MMR proteins, including *MutS* and *PMS1*, destabilize mismatched recombination intermediates to favor homologous over homeologous pairing. Finally, spindle checkpoint components, such as *MPS1-2*, ensure accurate chromosome orientation and segregation during meiosis.

### Barriers to autopolyploids as progenitors of allopolyploids

The limited contribution of autopolyploids to the origin of allopolyploids raises questions about their suitability as allopolyploid progenitors. Despite harboring alleles pre-adapted to polyploidy, we propose that autopolyploids may be constrained in two key ways. First, autopolyploids with four homologous chromosome copies can accumulate deleterious mutations due to relaxed purifying selection(Baduel et al., 2019; Bao et al., 2022). When an autopolyploid gamete contributes to the origin of an allopolyploid, these mutations may become exposed and, if not fully compensated by a homeolog, could lead to reduced fitness and increased extinction risk. Second, autopolyploids like *A. arenosa* exhibit altered meiotic crossover patterns(Bomblies et al., 2016). Whether these altered crossover patterns are beneficial or reversible or could be disruptive for homeolog segregation in allopolyploids remains unknown, but it may further reduce their viability as allopolyploid parents.

### Homeolog compensation and early stages of rediploidization

We find evidence of relaxed purifying selection in *A. suecica*, consistent with the idea that polyploidy confers increased mutational robustness through genetic redundancy. Previous studies have shown that diploid *A. thaliana* is more sensitive to mutagenesis than both natural and synthetic *A. suecica*(Spoelhof et al., 2021), supporting the claim that *A. suecica* has enhanced mutational robustness compared to its diploid parents. Despite the evidence of relaxed purifying selection, *A. suecica* has accumulated relatively few genes with a *de novo* LoF mutation suggesting that selection has still remained sufficiently strong to limit the accumulation of new LoF variants.

We observed bias in *A*. *suecica* toward LoF mutations in the *A*. *arenosa* sub-genome, however, most of these LoF mutations also segregate in *A. arenosa* and are likely inherited. This higher number of inherited mutations from the *A. arenosa* sub-genome are due to the higher genetic diversity and outcrossing nature of *A. arenosa*, which allows it to better mask deleterious mutations compared to the highly homozygous and predominantly selfing *A. thaliana*. *De novo* LoF mutations did not show sub-genome bias, consistent with our previous study, showing absence of sub-genome dominance on the transcriptional level(Burns et al., 2021).

Our analysis of LoF mutations in *A. suecica* reveals a pattern of compensation between homeologs, whereby the retention of at least one functional copy mitigates the impact of deleterious mutations. Specifically, when a LoF mutation arises in one homeolog, the other tends to remain intact more often than expected by chance. We find that these retained homeologs are under purifying selection, suggesting a selective pressure to preserve essential gene functions. This pattern aligns with findings in yeast, where duplicate genes can enhance mutational robustness but are often rapidly inactivated by LoF mutations, rather than undergoing gradual subfunctionalization(Mihajlovic et al., 2024). The homeolog compensation pattern is symmetric across sub-genomes; we find no bias in the accumulation of *de novo* LoF mutations or in the representation of dosage-sensitive single-copy genes. These patterns suggest that, shortly after allopolyploidization, mutations occur at random across sub-genomes, with selection acting *post hoc* to retain at least one functional copy per gene pair. While relaxed purifying selection allows LoF mutations to accrue, the compensatory retention of one intact copy buffers against gene function loss. This buffering, leveraging both standing and *de novo* variation, likely facilitates early genome stabilization in polyploids and helps maintain overall functional integrity. Yet this same genetic robustness may come at a cost: by masking the effects of deleterious mutations, homeolog compensation facilitates their persistence, setting the stage for gene loss to occur over time. We propose that homeolog compensatory dynamics mark the very beginning of rediploidization, a process by which the initial robustness of a polyploid genome ultimately shapes its return to a diploid-like state.

## Supplemental Figures

**Supplementary Figure 1:**
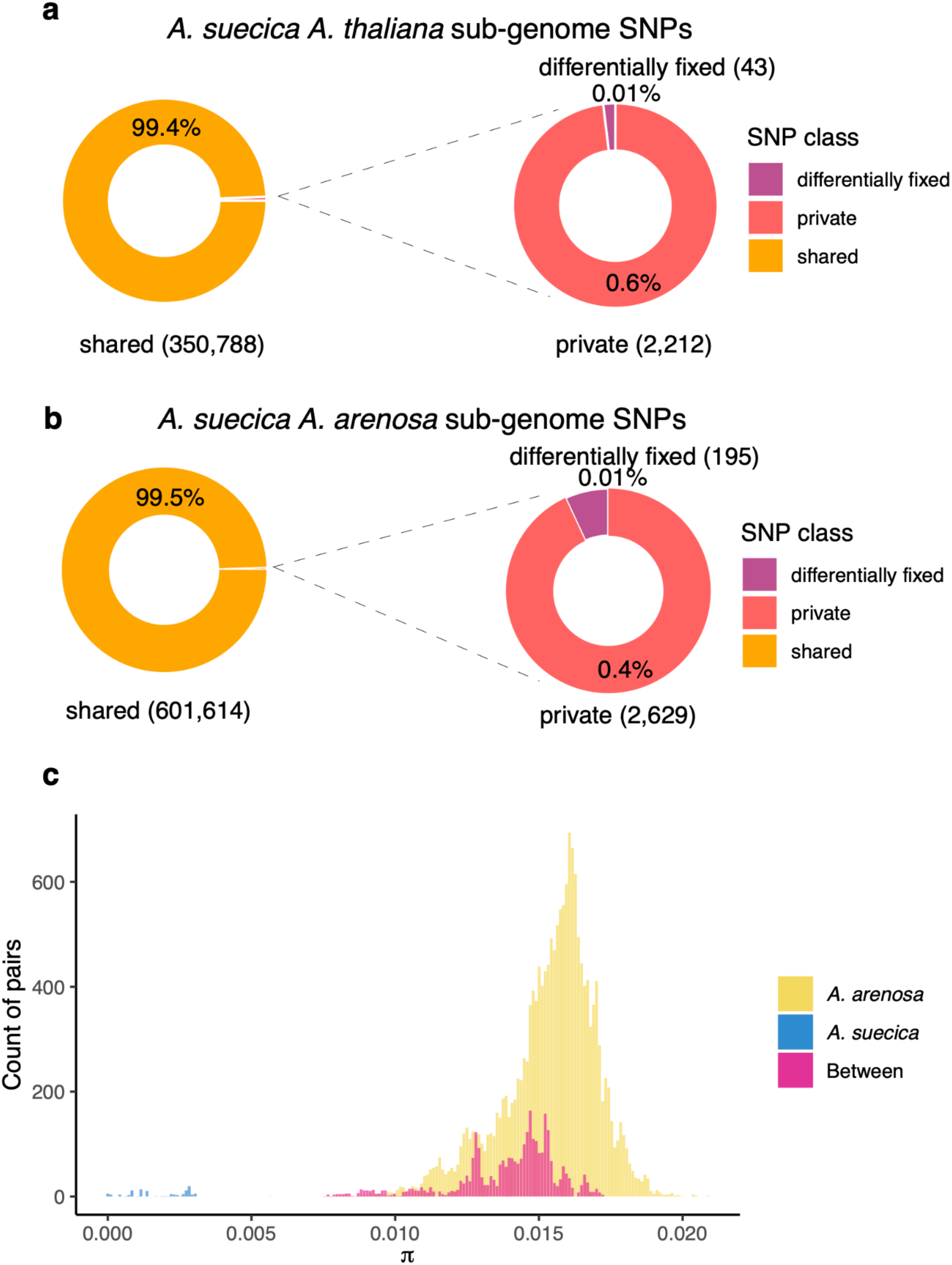
SNP Sharing and Nucleotide Diversity (π) in *A. suecica*. **a, b** Over 99% of *A. suecica* SNPs are shared with progenitor species (*A. thaliana* and *A. arenosa*), indicating contributions from multiple individuals of both progenitors. **c** Nucleotide diversity (π) is significantly lower in *A. suecica* (blue) compared to *A. arenosa* (yellow) due to the allopolyploidy bottleneck and self-compatibility. In contrast, the obligate outcrosser *A. arenosa* displays greater diversity within species than between species (pink).

**Supplementary Figure 2:**
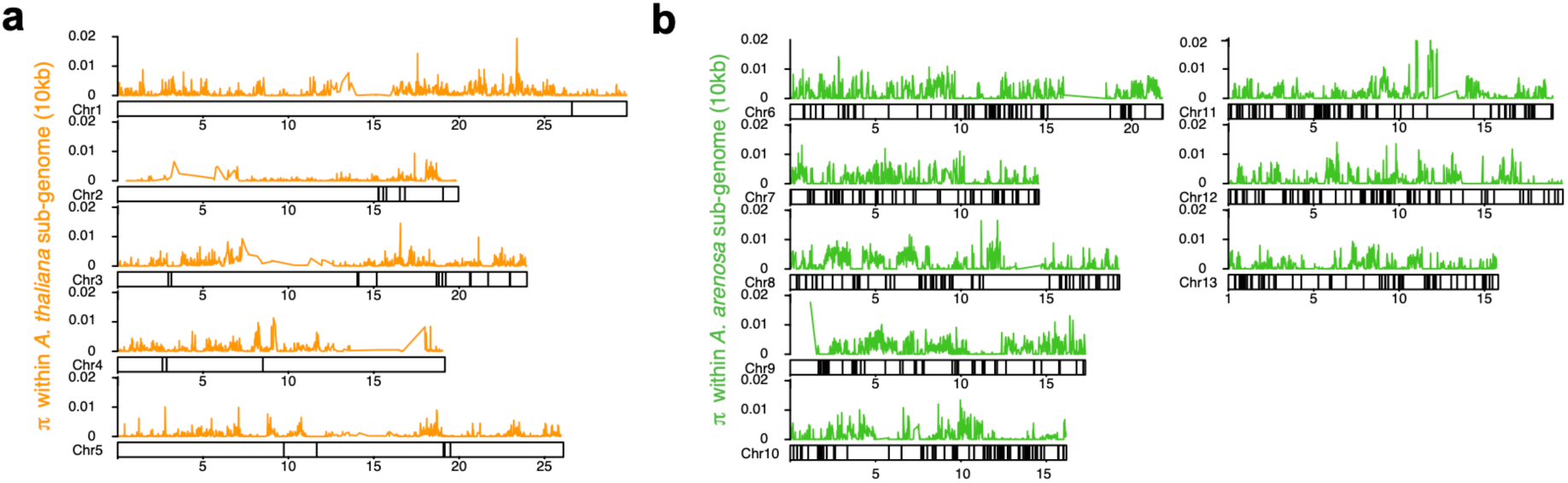
**a** In the *A. thaliana* sub-genome (Chr1 to 5), 35 genes, and **b** in the *A. arenosa* sub-genome (Chr 6 to 13), 585 genes show signatures of positive selection. These genes are found outside of the areas of low nucleotide diversity (π) within *A. suecica*, meaning a founder effect or population bottleneck is not the likely explanation for the signature of positive selection.

**Supplementary Figure 3:**
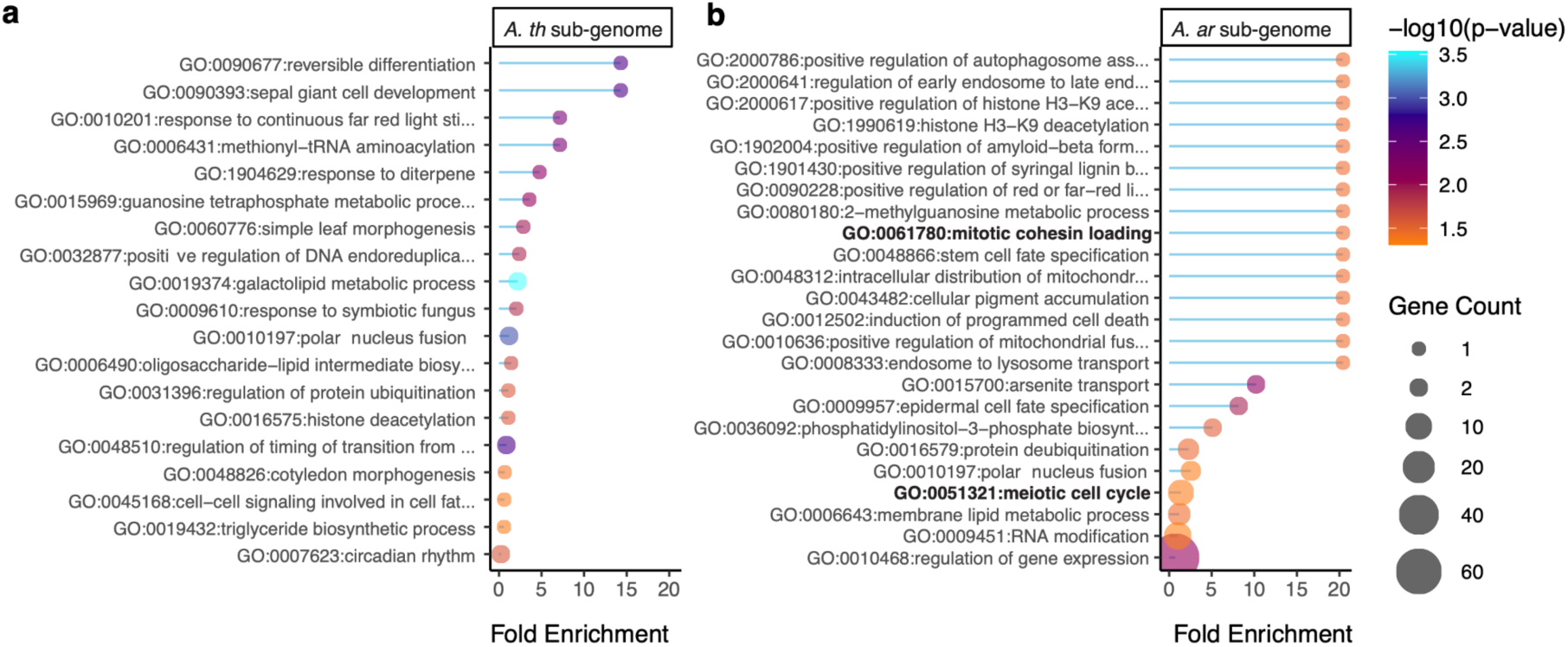
GO analysis of selection scan genes in *A. suecica*. GO enrichment for the **a** 35 genes on the A. thaliana sub-genome and **b** the 585 genes on the A. arenosa sub-genome. GO terms mentioned in the text are highlighted in bold.

**Supplementary Figure 4:**
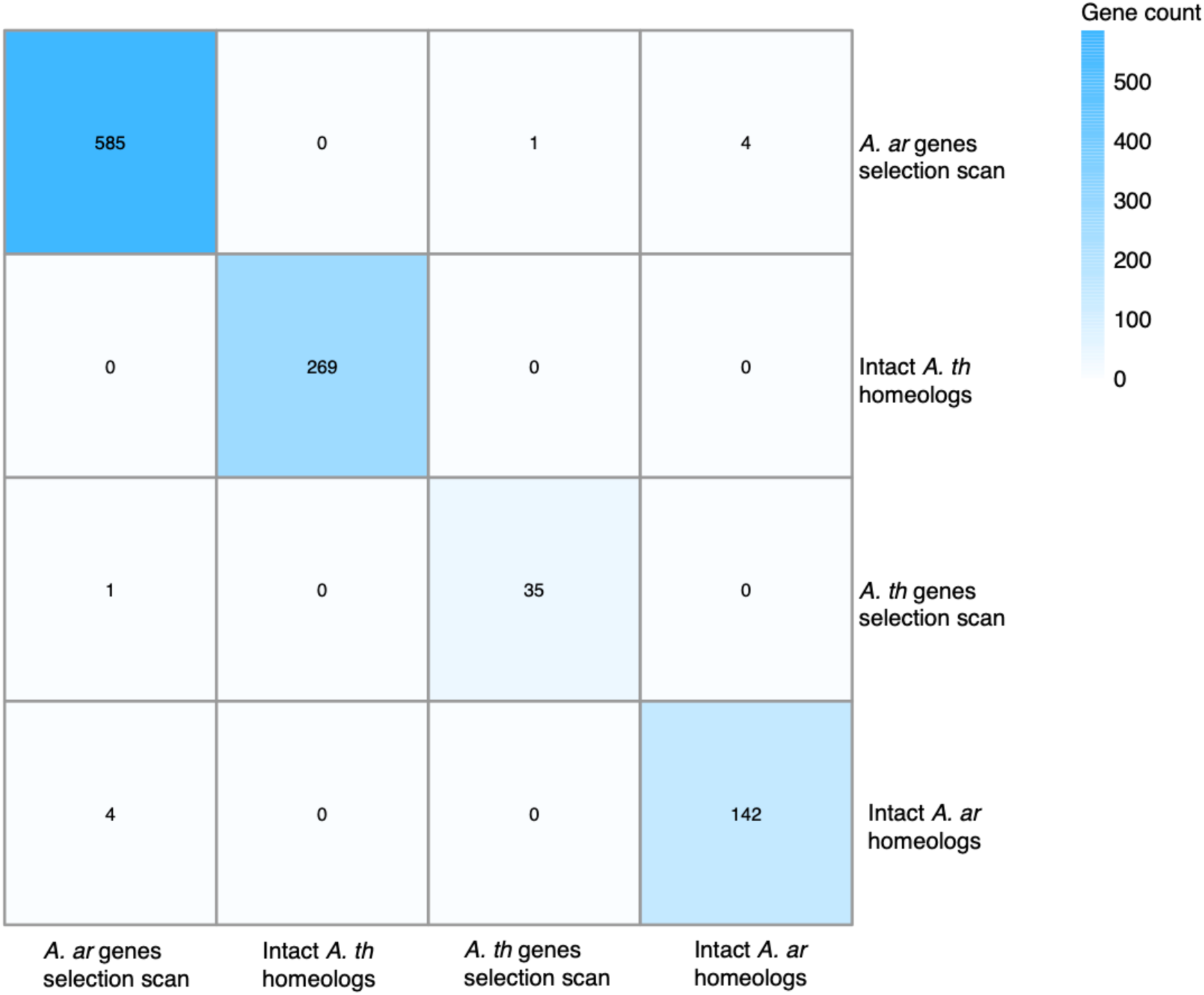
Overlap of genes in selection scans and homeologous gene pairs carrying one LoF mutation. Minimal overlap was observed between genes from selection scans and homeologous gene pairs with one intact copy and one carrying a LoF mutation, indicating that LoF on one homeolog is not driving the likely adaptive genetic changes on the other.

**Supplementary Figure 5:**
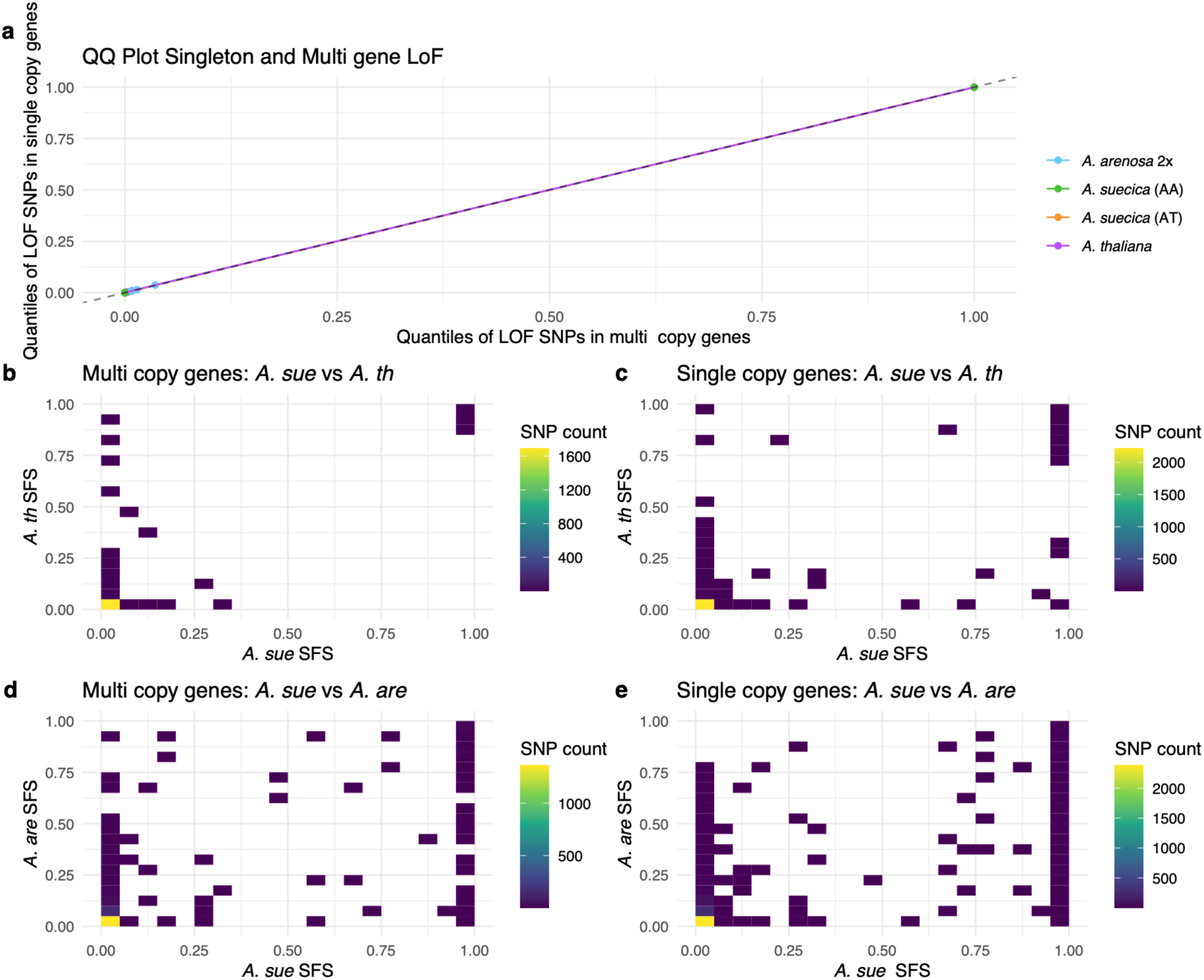
Lack of enrichment for LoF mutations in single-copy genes in *A. suecica.* **a** QQ plots comparing the population frequencies of LoF SNPs in single-copy genes versus multi-copy genes in *A. suecica*. No significant increase in the population frequency of LoF mutations is observed in *A. suecica* for single-copy genes, indicating that both single- and multi-copy genes are under similar selective pressures. **b-e** Joint allele frequency spectra for LoF SNPs, comparing *A. suecica* and the parental species for multi-copy and single-copy genes. The data show an enrichment of singleton SNPs for all species and gene copy status, consistent with strong purifying selection acting on these deleterious mutations.

**Supplementary Figure 6:**
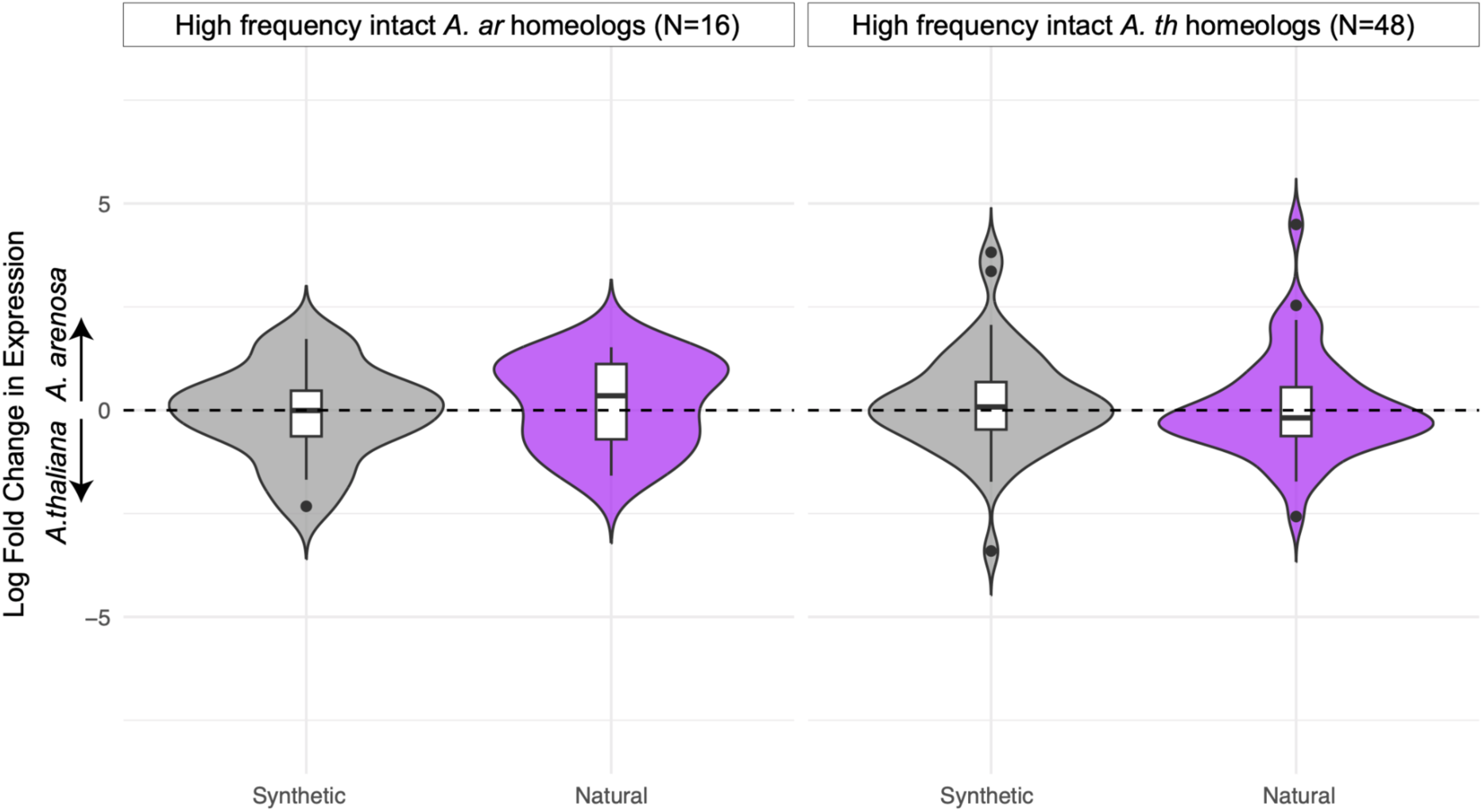
Sub-genome expression bias towards intact homeolog. Violin plots show the log_2_ fold change in gene expression between homeologs in synthetic versus natural *A. suecica* accessions. Expression bias is plotted for genes where one homeolog exhibits a high-frequency loss-of-function mutation (≥60% in the population) while the homeolog either on the *A. arenosa* sub-genome (left panel, N = 16) or in the *A. thaliana* sub-genome (right panel, N = 48) remains intact. Each violin represents the distribution of expression bias for either synthetic (gray) or natural (purple) *A. suecica*. A log fold change of zero (dashed line) indicates balanced expression between homeologs. Expression in natural *A. suecica* is biased toward the intact homeolog, consistent with purifying selection maintaining expression of the functional copy. This expression bias is absent in synthetic *A. suecica*, indicating that it is not an inherited regulatory pattern but rather an evolved expression change due to the presence of a nonfunctional allele in the other homeolog.

**Supplementary Figure 7:**
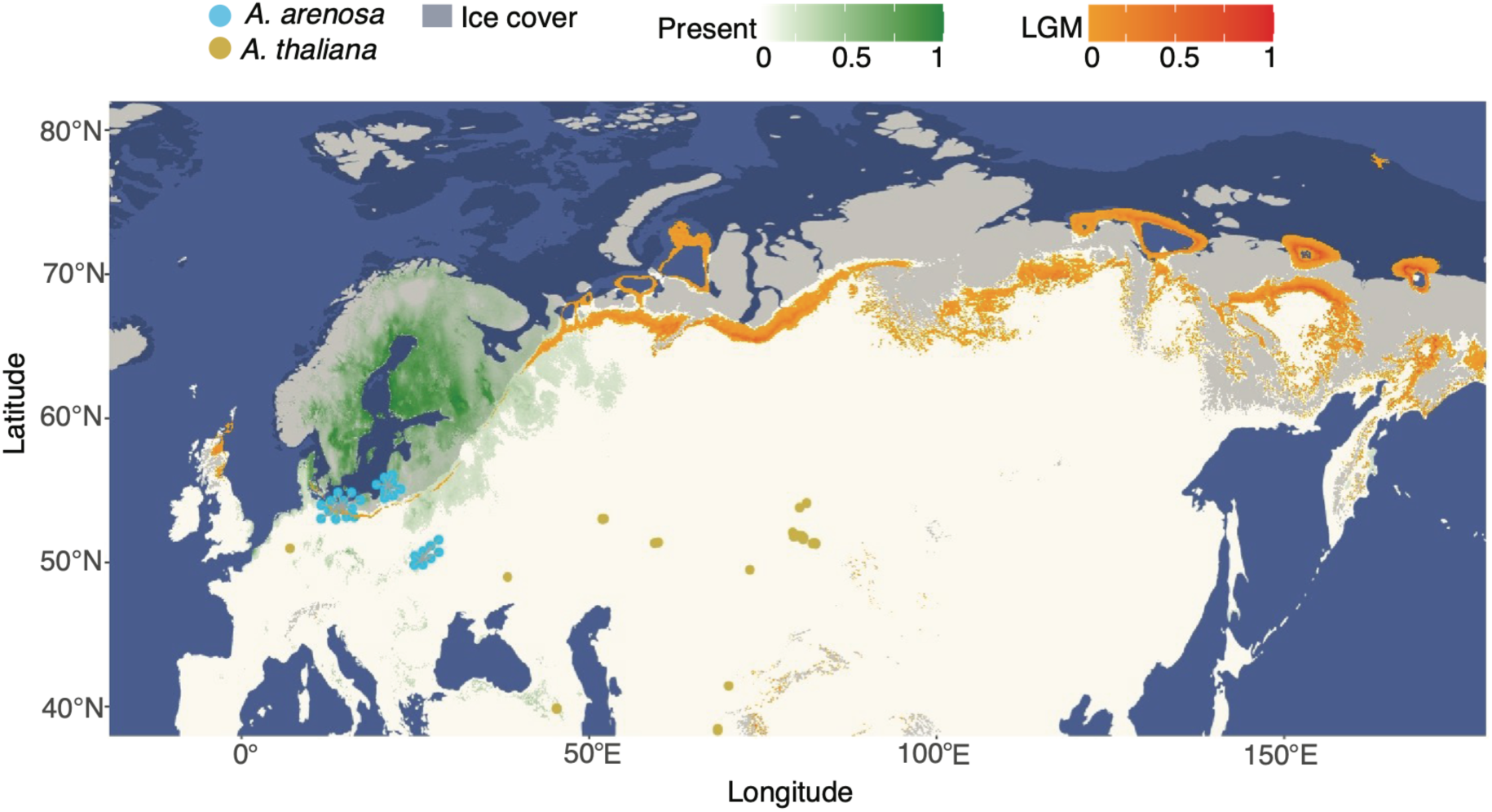
The geographic origin of *A. suecica*. Niche modeling of *A. suecica* shows probability of occurrence for the current climate (green) and the LGM (orange). Closest parental populations include Southern Baltic and Ukrainian diploid *A. arenosa*, and the top 5% of *A. thaliana* accessions genetically closest to *A. suecica*.

## Code availability

Custom scripts used in this study are available at https://github.com/burnsro/ArabidopsisSuecica/tree/main/origin

## Data availability

Raw fastq files of the Illumina sequenced herbarium samples for 5 tetraploid *A. arenosa* and 1 *A. suecica* generated in this study were deposited to the European Nucleotide Archive (ENA) under the project number PRJEB79527.

## Author contributions

R.B and P.Y.N conceived the study. Herbarium samples were aquired by P.Y.N and genomic DNA was extracted by R.B. R.B, P.Y.N, A.K, A.G, U.K.K, A.D.S analyzed the data. R.B, P.Y.N, F.K and A.D.S wrote the paper.

## Acknowledgements

This work was supported by an EMBO long-term postdoctoral fellowship ALTF224-2022 to R.B. P.Y.N. acknowledges the European Union (ERC, HOW2DOUBLE, 101041354). The views and opinions expressed are, however, those of the author(s) only and do not necessarily reflect those of the European Union or the European Research Council Executive Agency. Neither the European Union nor the granting authority can be held responsible for them. F.K. and P.Y.N. acknowledge the joint Deutsche Forschungsgemeinschaft (DFG) and Czech Science Foundation (GACR) program, project number 490698526 and 22-29078K. F.K. and P.Y.N. also acknowledge the DAAD exchange travel grant, project number 57601834. We thank Heïdi Serra, Andrew Lloyd and Nicola Gorringe for helpful comments on the manuscript and discussions.

